# Functional characterization of red-shifted rhodopsin channels from giant viruses explored by a machine-learning model for long-wavelength optogenetics

**DOI:** 10.1101/2025.09.16.676488

**Authors:** Shunki Takaramoto, Chenxiang Zhao, Masayuki Karasuyama, Frederik Schulz, Masaya Watanabe, Yuma Kawasaki, Takashi Nagata, Masae Konno, Naoya Morimoto, Masahiro Fukuda, Yu Inatsu, Hiromu Yawo, Oded Béjà, Hideaki E. Kato, Tanja Woyke, Ichiro Takeuchi, Keiichi Inoue

## Abstract

Channelrhodopsins (ChRs) are light-gated ion channels. These proteins are widely used in optogenetics to optically manipulate neural activity. However, manipulation using short-wavelength light to activate ChRs causes cell toxicity and is hampered by low tissue penetration. To overcome these difficulties, although several red-shifted ChR variants have been identified, further red-shift is required for more efficient and noninvasive neural control. While molecular screening of ChRs requires high-cost experiments, recent machine-learning-based protein functionality prediction enables more efficient selection of target proteins for characterization. Here, we constructed an *elastic-net* machine-learning model trained on 1,163 experimental data to predict the maximum absorption wavelength (*λ*_max_) of uncharacterized ChRs. The model suggested several red-shifted candidates, and we identified a viral rhodopsin channel, ChR024, with the second-longest *λ*_max_ as a cation-conducting ChR (*λ*_max_ = ∼578 nm) after Chrimson (*λ*_max_ = ∼580 nm). This result demonstrates the high impact of ML on reducing the screening costs of functional proteins.

## Introduction

Channelrhodopsins (ChRs) are sub-class of microbial rhodopsins functioning as light-gated ion channels using an all-*trans*-retinal chromophore^1^. Cation and anion-specific ChRs, CCR and ACR, are found in the genomes of many eukaryotic microorganisms and giant viruses^2^. CCR and ACR heterologously expressed in animal neurons are capable of inducing or suppressing the neuronal firing by destabilizing and hyperpolarizing the neurons in a light-dependent manner, respectively. Hence both CCR and ACR are widely used in optogenetics to understand the causal relationship between the respective neuronal circuits in the animal body and phenotypes such as locomotion, memory, emotion, social behavior, etc^3^. To enable more non-invasive and versatile optogenetic manipulation, ChRs that can be manipulated with longer-wavelength light have been explored and developed by gene mining and protein engineering^4–7^, because the wavelength of light is longer, the photo-toxicity for the cells and the penetration depth in animal tissue are lower and deeper, respectively.

However, the hit rate of the screening for new red-shifted ChRs is not high, leading to increased experimental costs. To reduce the cost associated with gene mining and protein engineering for the exploration and development of highly red-shifted ChRs, machine-learning (ML) models are valuable for predicting the absorption maximum wavelengths (*λ*_max_) of the retinal chromophore in rhodopsins. Such models can help identify promising proteins with red-shifted absorption without screening experiments^8–12^. Here we applied an ML model trained on 1,163 experimental data collected from the literature and from datasets constructed by some of the authors.^8,9^ This model was then used to predict the *λ*_max_ of 259 new ChR homologes identified through homology searches of the public gene databases, based on their amino acid residues surrounding the retinal chromophore.

## Results

### Prediction and experimental measurement of the *λ*_max_ of ChRs

We trained an ML model that predicts the *λ*_max_ of ChRs from their amino-acid sequences, using 1,163 training instances, each containing information on the amino acid sequence and the experimentally determined wavelength^8,9^. As the choice of ML model, we employed a well-known sparse linear model called the *elastic-net model*. Because some of the authors previously found that 24 residues surrounding the retinal chromophore play a critical role in determining the *λ*_max_ of microbial rhodopsins (Supplementary Fig. 1)^9^, the ML model construction and the *λ*_max_ prediction were conducted based on these residues (Supplementary Table 1). In the employed ML model, 18 physico-chemical features of these 24 amino acid residues were used as features (Supplementary Data 2 in Ref.^9^), i.e., the sparse linear model was trained with a total of 432-dimensional features.

The *λ*_max_ of 259 ChR-like proteins were predicted using this ML model, and 23 genes predicted to have the most red-shifted *λ*_max_ were synthesized and transfected into mammalian (COS-1) cells (Fig. 1a). The cells expressing 8 out of 23 rhodopsin genes exhibited strong orange, red, or purple colors. The *λ*_max_ of these rhodopsins were determined by measuring the UV-visible absorption change upon the hydrolysis of the retinal Schiff-base (RSB) linkage by hydroxylamine (HA, Fig. 1b)^13^. Notably, ChR024, which was named based on the serial number in the list of 259 ChR homologue genes, exhibited the most red-shifted *λ*_max_ at 578 nm (Fig. 2a). When ChR024 C-terminally labeled with fluorescence protein (eYFP) was expressed in rodent ND7/23 cells, the expressed protein was well localized on the plasma membrane (Fig. 2b). The light-dependent ion-transport activity of ChR024 was measured by patch clamp experiments using bath and pipette solutions containing 142 and 132 mM Na^+^, respectively. Upon 575-nm light illumination, photocurrents that reversed direction in accordance with changes in membrane potentials were observed (Fig. 2c). The reversal membrane potential (*E*_rev_) with 142- and 132-mM Na^+^ bath and pipette solutions was 11 ± 2 mV. Compared with *E*_rev_ obtained using 146 mM N-methyl-D-glucamine (NMG) bath and 130 mM NMG pipette solutions, a significant shift in *E*_rev_ was observed with 146 mM Na^+^ or Li^+^ bath solutions (Fig. 2d and Supplementary Fig. 2) indicating ChR024 is a CCR absorbing 578-nm yellow light. This is natural CCR with the second longest *λ*_max_ after Chrimson (*λ*_max_ = ∼580 nm)^14^.

**Fig. 1.**
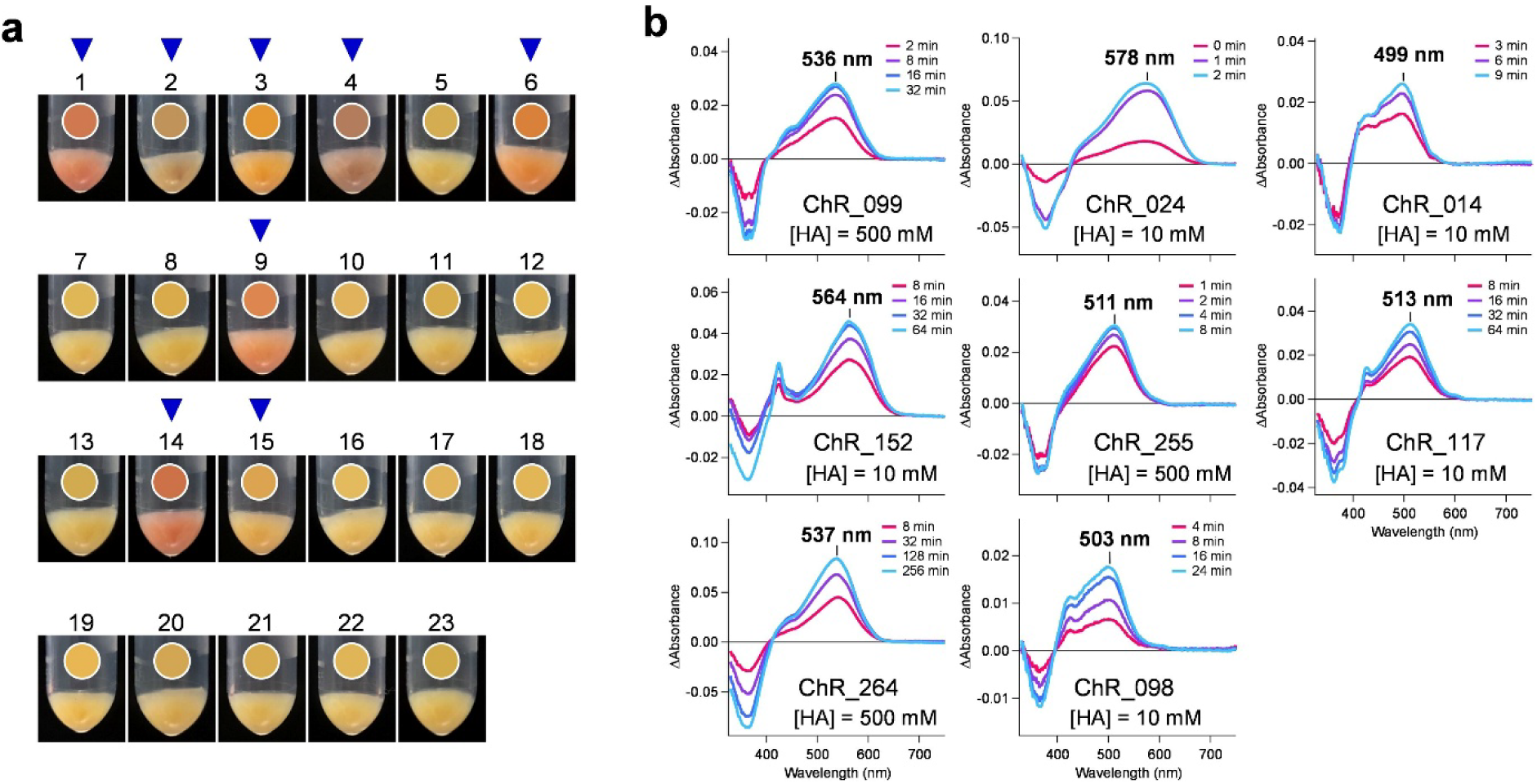
*λ*_max_ value analysis of ChRs predicted to exhibit red-shifted absorption. **a** Photographs of pellets of COS-1 cells transfected with ChR genes that were predicted to exhibit red-shifted absorption based on ML calculation. Pixel colors at the center of each pellet are shown in circles above the corresponding samples. Samples displaying pronounced rhodopsin coloration and subjected to *λ*_max_ measurements in panel b are indicated by arrowheads. **b** Difference UV-visible absorption spectra of rhodopsins in solubilized membrane, recorded before and after photobleaching via hydrolysis of RSB linkage in the presence of HA. Spectra corresponding to different illumination durations are shown in different colors.

**Fig. 2.**
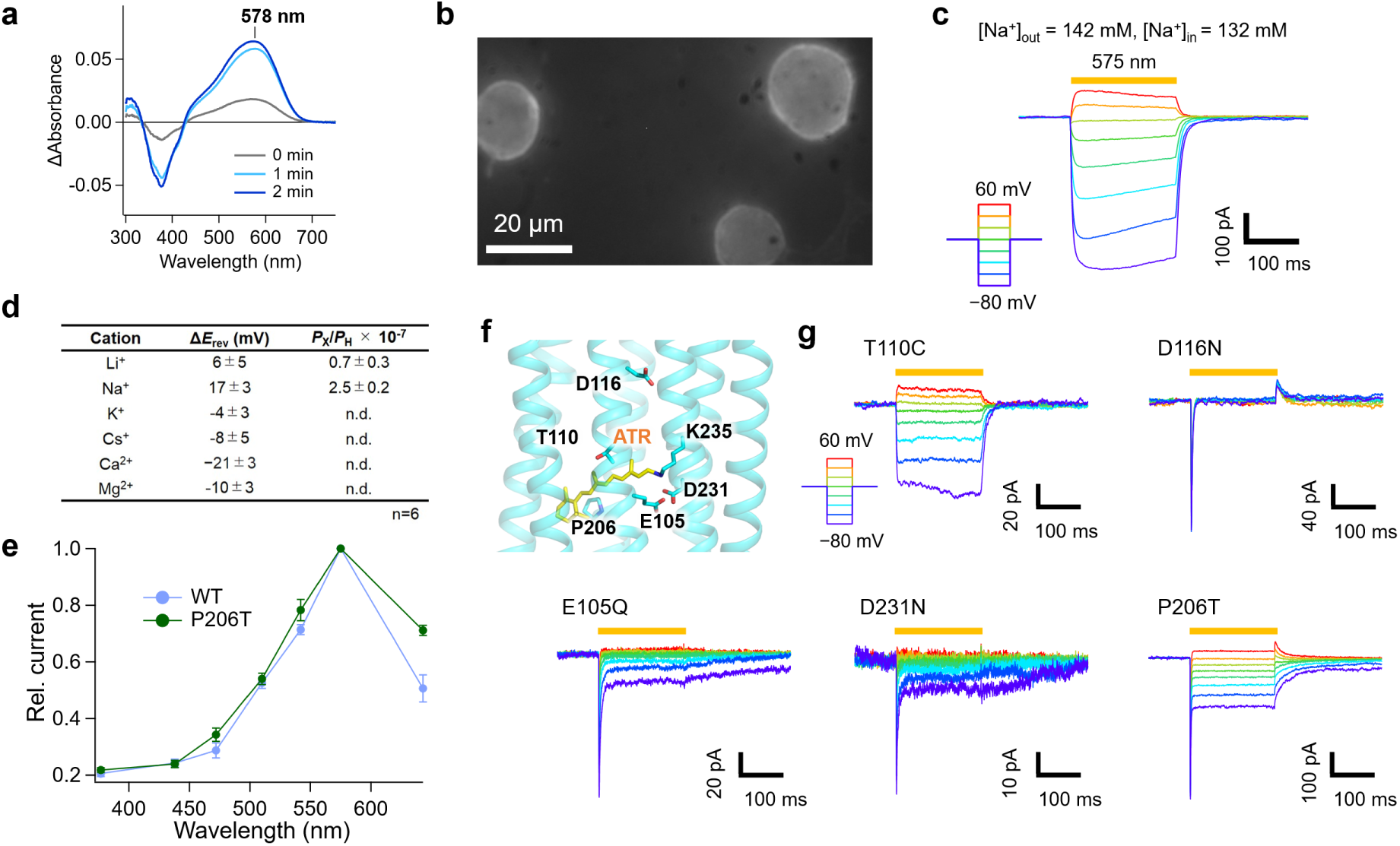
Exploration of red-shifted ChRs and electrophysiological characterization. **a** Difference UV-visible absorption spectra of ChR024 in solubilized membrane of COS-1 cells, recorded before and after photobleaching via hydrolysis of RSB linkage with HA. Spectra with different illumination periods are shown in different colors. **b** Fluorescence image of ND7/23 cells expressing ChR024 C-terminally labeled with eYFP (Scale bar = 20 μm). **c** Photocurrents of ChR024 at 142 mM [Na^+^]_out_ and 132 mM [Na^+^]_in_ at pH = 7.4. Yellow line: period of light illumination at 575 nm. **d** Difference in *E*_rev_ between different cations and H^+^ (Δ*E*_rev_) and relative permeability of different cations against H^+^ (*P*_X_/*P*_H_) (mean ± S.E., *n* = 4– 5). **e** Normalized action spectra of the WT and P206T. **f** Schematic illustration of mutated positions based on a structure modeled by Alphafold2. **g** Photocurrents of ChR024 mutants at 142 mM [Na^+^]_out_ and 132 mM [Na^+^]_in_ at pH = 7.4. Yellow line: period of light illumination at 575 nm.

### Electrophysiological study on the ion-transport mechanism of ChR024

To investigate the ion-transport mechanism of ChR024, we compared its amino acid sequence with other CCR and ACR (Supplementary Fig. 3). The amino acid sequence of ChR024 is highly different from those of other CCRs and ACRs (Supplementary Fig. 3) and it is located in a branch distinct from the branches of other CCR/ACR in the phylogenetic tree of microbial rhodopsins (Supplementary Fig. 4). Notably, a cysteine residue highly conserved among CCR and ACR (e.g. C128 in *Cr*ChR2, Supplementary Fig. 1) in the third transmembrane helix (TM3) and forming the so-called DC gate with aspartic acid (D156 in *Cr*ChR2) in the fourth transmembrane helix (TM4) is substituted with threonine (T110) in ChR024, indicating the ion-transport mechanisms of ChR024 differs substantially from those of other CCRs.

If T110 is replaced with cysteine, the photocurrent was significantly reduced compared to the wildtype (WT) ChR024 (Figs. 2f and g). The structure of ChR024 modelled using AlophaFold2^15^ suggested that two acidic residues, E105 and D231, are located on the extracellular side of the Schiff base linkage (–C=N–) of the retinal (Fig. 2f). The acidic residues at these positions are generally deprotonated in most microbial rhodopsins and function as counterions stabilizing the protonated state of the RSB^16^. The mutations of these residues, E105Q and D231N, exhibited highly reduced photocurrents compared to the WT (Fig. 2g), indicating they play a critical role in the cation transport. Additionally, there is an aspartic acid, D116, on the cytoplasmic side at the position homologous to D96 bacteriorhodopsin (BR) (Fig. 2f and Supplementary Fig. 3). Whereas D96 is known to work as a proton donor during the photocycle of BR, the role of D116 in ChR024 is not clear. The mutation of D116 to asparagine (ChR024 D116N) resulted in the complete loss of channel currents (Fig. 2g). Hence, D116 is critical to the ion-channel function of ChR024, which also differs the ion-transport mechanism of canonical CCRs.

In the sixth transmembrane helix (TM6), there is a proline residue, P206, near the β-ionone ring of the retinal (Fig. 2f). While a homologous proline residue is highly conserved in many microbial rhodopsins (Supplementary Fig. 3), the mutation of this proline to threonine is known to systematically red-shift the *λ*_max_ of the retinal^13^. To further red-shift the absorption of ChR024, ChR024 P206T mutant was constructed. Notably, the action spectrum of ChR024 P206T exhibited higher sensitivity to red light than that of the WT (Fig. 2e), indicating the P-to-T mutation is effective to red-shift the absorption of ChR024 as other microbial rhodopsins, and ChR024 P206T would be useful for optogenetic application using longer-wavelength light.

### Spectroscopic study on the ion-transport mechanism of ChR024

To spectroscopically study the ion-transport mechanism of ChR024, the protein was expressed in *Spodoptera frugiperda* (Sf9) insect cells using the pFastBac baculovirus system. Purified protein by Ni-NTA affinity and gel-filtration chromatography exhibited a broad peak (FWHM = 138 nm) at 576 nm (Fig. 3a). This is consistent with the spectrum obtained by HA bleaching method (Fig. 2a) and the action spectrum (Fig. 2e). The *λ*_max_ was shifted from 579 to 520 nm by raising pH from 7.0 to 9.3 (Fig. 3b and c). Similar blue shift from the longer-wavelength broad peak was observed for Chrimson and the rhodopsin domains in bestrhodopsin, and it was suggested to be induced by the deprotonation of acidic residue near the RSB^14,17^, indicating that either of the counterion residues, E105 or D231, is protonated at physiological pH and essential for the red-shifted absorption of ChR024.

**Fig. 3.**
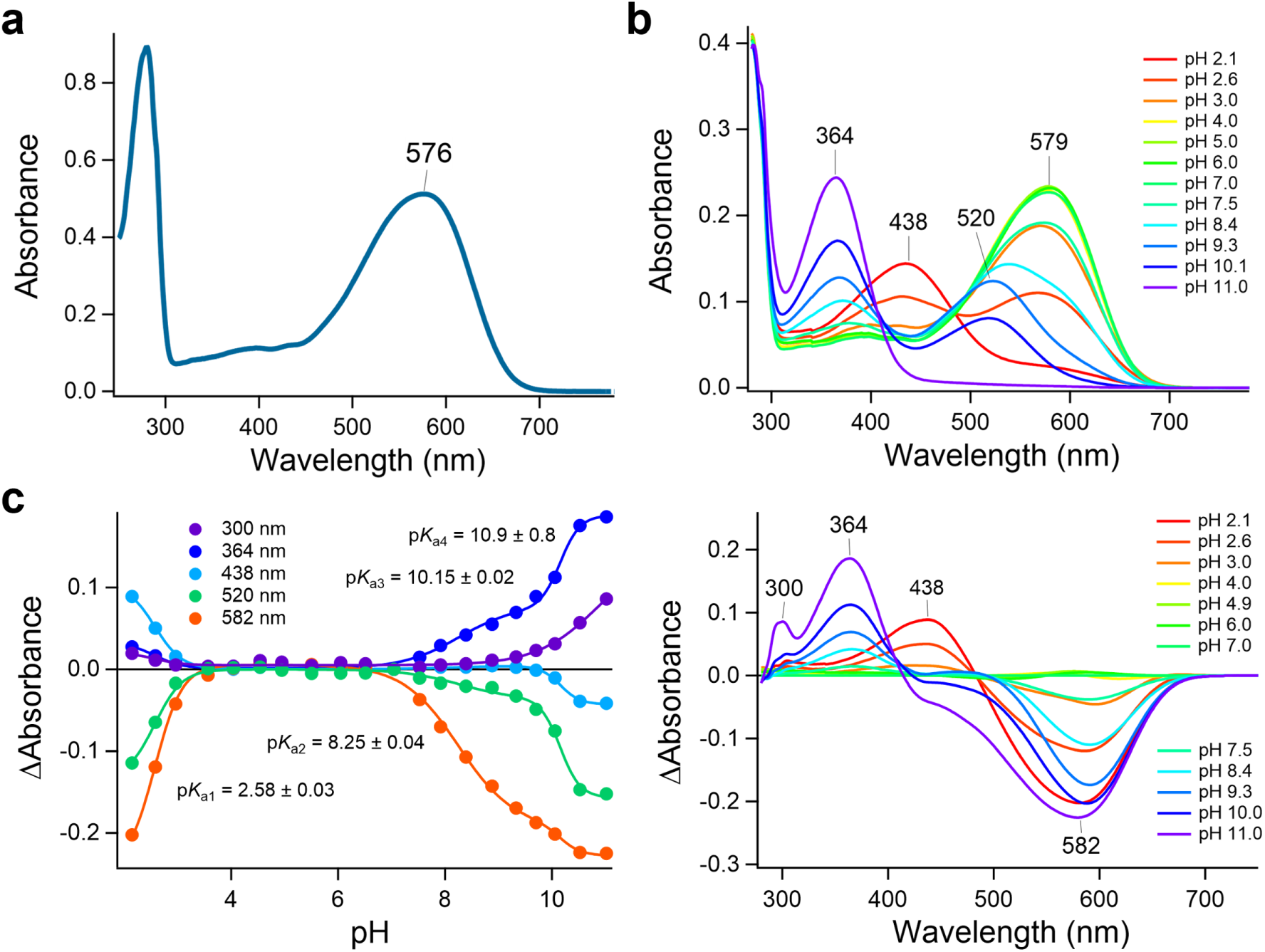
Spectroscopic characterization of purified ChR024. **a** The absorption spectrum of purified ChR024 in 20 mM HEPES-NaOH (pH 7.5), 100 mM NaCl, 0.03% DDM. **b** The pH dependence of the absorption spectra (to) and the difference absorption spectra calculated by subtracting the spectrum at pH 7 from the one at each pH (bottom). **c** Absorption change at specific wavelengths. All data were fitted by the quadruple Henderson–Hasselbalch equation, and four p*K*_a_ values were shown in the graph.

To investigate the photo-reaction and channel opening/closing dynamics of ChR024, transient absorption measurements of purified protein in the UV-visible region and the patch clamp measurement using a nanosecond pulsed laser were performed (Fig. 4a and b). Because a decrease in all-*trans*-retinal upon illumination was observed by the high-performance liquid chromatography (HPLC) of retinal oxime produced by the hydrolysis reaction of the RSB with HA, together with an increase in the proportion of 13-*cis*-retinal (Supplementary Fig. 5), ChR024 binds all-*trans*-retinal in the dark, and it photo-isomerizes to the 13-*cis* form as most microbial rhodopsins^16^. Then, the blue-shifted photointermediates similar to the L, M, and N intermediates observed in the photocycle of BR^16^ appeared in the microsecond to millisecond time region (Figs. 4a–c). Although the most blue-shifted M intermediate represents the deprotonated state of the RSB, its accumulation is extremely small, and the RSB is thought to be mostly kept protonated throughout the entire photocycle of ChR024. Then, the channel opening and closing occurred at 2.0 and 4.3 ms after the photo-excitation with a minor closing component at 22.5 ms (Fig. 4b). If we compare these time constants with the lifetimes of the photointermediates determined based on the transient absorption change, the channel opening occurs on the N formation, and it is spectrally silently closed at 4.3 ms. Hence, the conformational change associated with the channel closing does not affect the structure of the RSB.

**Fig. 4.**
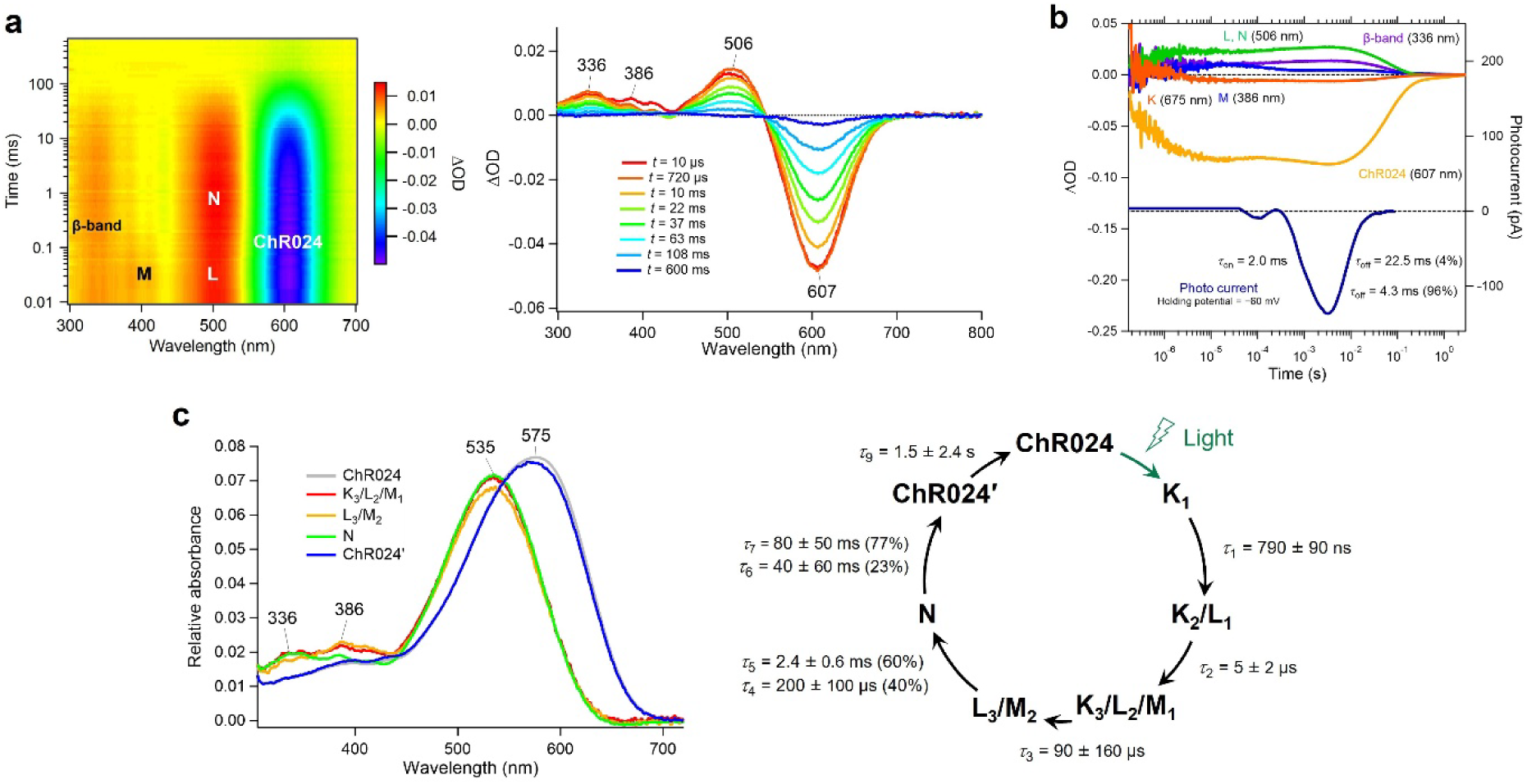
Laser-flash photolysis analyses of the photocycle of ChR024. **a** Two-dimensional plot of the transient absorption change of ChR024 (left) and the transient absorption signal at specific time points (right). **b** Time courses of the transient absorption change at specific wavelengths (upper) and photocurrent after 5-ns laser pulse illumination (lower). **c** Absolute absorption spectra of the initial state and photointermediates (left) and the photocycle model of ChR024 (right) based on the multi-exponential global analysis of the transient absorption change in **a** and **b**.

In contrast to other CCRs generally exhibiting a strong peak current at the onset of light, no peak current was observed for ChR024 upon 200-ms illumination (Fig. 2c). However, we noticed that a peak current appeared only on the first illumination and disappeared on subsequent illuminations (Supplementary Figs. 6a and b). However, when we illuminate nanosecond pulses at 1-s interval, the photocurrent intensity gradually recovered (Supplementary Fig. 6c), and a peak current again appeared during subsequent prolonged illumination (Supplementary Fig. 6d). If we investigate the isomeric state of the retinal, the protein after prolonged illumination contained higher amounts of 13-*cis* and 11-*cis* retinal than that in the dark state (Supplementary Fig. 5). However, interestingly, if the protein is exposed to nanosecond laser pulses after the prolonged illumination, the proportion of all-*trans* form was recovered. These results indicate that, while prolonged illumination converts ChR024 to a less functional state with 13-*cis*/11-*cis* forms through the photo-excitation of a photo-intermediate(s). This less functional state is reverted to the original all-*trans* form by nanosecond laser pulses, and the recovered state once again exhibits strong peak currents.

### Rhodopsin genes located near the ChR024 gene in the genomes

Notably, we found additional rhodopsin-like genes downstream of ChR024 and its homologue genes in the original scaffolds (green arrows in Supplementary Fig. 7a). One of them, which has a unique serine–serine–asparagine (SSN) motif at the positions corresponding to D85– T89–D96 in BR^18^, could be expressed in COS-1 cells and exhibited pinkish color (Supplementary Fig. 7b), indicating this rhodopsin can incorporate retinal in its protein body with a *λ*_max_ at 530 nm (Supplementary Fig. 7c). Notably, Ser74 and Asn194 at the positions corresponding to ChR024 E105 and D231 in TM3 and TM4, respectively, suggest that the RSB is stably protonated without typical counterions. However, it exhibits no photocurrent in ND7/23 cells, indicating its biological function is different from light-dependent ion transport.

## Discussion

Long-wavelength-absorbing ChRs are in demand for optogenetic manipulation using red or near-infrared light. However, to identify or to develop red-shifted ChRs, a large amount of experimental screening is necessary, imposing high costs. Regarding this problem, ML-based prediction of the *λ*_max_ of uncharacterized rhodopsins can significantly improve the efficiency of the molecular screening and reduce experimental costs^8,9^. In this study, we constructed a new ML model, which employs *elastic-net model* and was trained with 1,163 experimental data of the *λ*_max_ of microbial rhodopsins and their variants, and predicted the *λ*_max_ values of uncharacterized ChR genes. Among them, the ChR024 exhibited the longest *λ*_max_ at 575 nm, which is close to the currently most red-shifted ChR Chrimson (∼580 nm).

The pH dependence of the absorption spectra of ChR024 indicated that the protonation of the counterion of the RSB significantly contributes to the red-shifted absorption of ChR024, which is similar to that known for Chrimson and other red-shifted microbial rhodopsins^14,17,19–21^.

While ChR024 can transport Na^+^ and Li^+^, the transport of larger cations (K^+^, Rb^+^, and Cs^+^) is negligible. Its proton selectivity (P_H+_/P_Na+_ = 14 × 10^6^) is similar to that of Chrimson (14 × 10^6^ ^19^) and higher than that of other CCRs (e.g., *Cr*ChR2: 1–2 × 10^6 22^; *Cr*ChR1: ∼1.0 × 10^6 23^; HulaChrimson: 1× 10^6 24^; *Ps*ChR: 0.4 × 10^6 19^; *Gt*CCR4: 0.02 × 10^6^ ^25^; and HulaCCR1 0.07 × 10^6 26^). Therefore, the counterion-protonated configuration observed in ChR024 and Chrimson is likely to facilitate H^+^ transport compared to that of other cations.

Interestingly, in the original scaffold, ChR024 gene is adjacent a gene coding a non-ion-transporting rhodopsin with an SSN motif (SSN rhodopsin). Although the function of SSN rhodopsin is unclear, it may modulate the function of ChR024 in a light-dependent manner, as known for a heterodimer of rhodopsin-guanylyl cyclase and its red-shifted homologue, neorhodopsin (NeoR)^20^.

After ChR024, ChR152 exhibits its *λ*_max_ at 564 nm (Fig. 1b). The phylogenetic analysis suggested that ChR152 is classified into the cryptophyte ACR clade, and its *λ*_max_ value is the fifth longest among 29 ACRs for which the *λ*_max_ or the peak wavelength of photocurrents has been reported. Therefore, CCR and ACR, with the second and the fifth longest *λ*_max_, respectively, were included in seven ChRs, which were predicted to have long *λ*_max_ by the ML model and expressed in mammalian cells, demonstrating the ML-based prediction of functional protein can significantly enhance the efficiency of molecular screening and reduce experimental costs. On the other hand, 16 out of 23 ChRs that were predicted to have long *λ*_max_ could not be expressed in mammalian cells, suggesting that the lack of the prediction of protein expression difficulties still prevents high-throughput screening with high hit rates. To overcome this difficulty, the acquisition of experimental datasets including not only *λ*_max_ values but also protein expression levels is necessary. Furthermore, combining an experimental design based on active learning with robotic molecular characterization^27^ would greatly accelerate the mining and development of functional proteins.

## Methods

### Construction of training and target data sets

In this study, we constructed a new training data set (Supplement Data Table 1) by adding 97 genes for which the *λ*_max_ had recently been reported in the literature or determined by our experiments, to a previously reported data set^8^. The sequences were aligned using ClustalW^28^, and the results were manually checked to avoid improper gaps and/or shifts in the TM parts. The aligned sequences were then used for ML-based modeling.

To collect microbial rhodopsin genes for the target data set, *Cr*ChR2 sequence was used as queries for searching homologous amino acid sequences in JGI IMG/M database, with the threshold E-value set at < 10 by default, and sequences with > 180 amino acid residues were collected. All sequences were aligned using ClustalW^28^, and the aligned sequences were used as the training data set for the prediction of *λ*_max_.

### ML modeling

Let us denote the training set of the ML model by 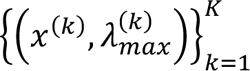, where *K* = 1,163 is the number of microbial rhodopsins in the database, *x*^(*k*)^ is the 432-dimensional feature vector which represents *M*(= 18) physico-chemical features of the *N*(= 24) amino-acid residues (Supplementary Table 1) of the *k*-th rhodopsin in the database, whereas *y*^(*k*)^ is the absorption maximum wavelength of the rhodopsin. As the choice of ML model, we employed the elastic net, which is formulated as

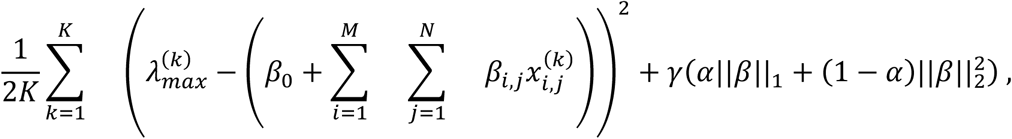

where β_0_ is a scalar parameter, β is a 432-dimensional parameter vector whose elements are {β_i,j_}, *i* = 1, …, *M*, *j* = 1, …, *N*. Furthermore, γ and α are the hyperparameters which govern the stability and sparseness of the trained ML models. The elastic-net is a well-known sparse linear model training algorithm, and we used a package called *glmnet* in the R programming environment. The elastic-net has two hyperparameters, γ > 0 and α ∈ [0,1]. When α = 1, the elastic-net reduces to the learning algorithm called Lasso, and when α = 0, it reduces to Ridge Regression. In this study, to maintain the sufficient sparsity of the model, we fixed α = 0.9 and selected γ through 10-fold cross-validation. After training the model, the maximum absorption wavelengths of a target rhodopsin is predicted as

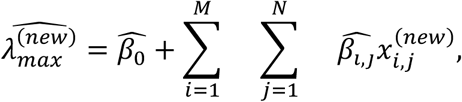

where 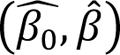 is the solution of the above optimization problem, 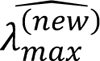 is represents the predicted maximum absorption wavelength of the target rhodopsin, and *x*^(new)^_i,j_ represents the j-th physico-chemical feature of the i-th residue of the target rhodopsin.

### The phylogeny of microbial rhodopsins

The 88 sequences of the highly conserved TM1–7 region of microbial rhodopsins were aligned by PROMALS3D multiple sequence and structure alignment server^29^. The 3-D structural data of following rhodopsins were used for the structural constraint: BR (PDB ID: 7Z09^30^), AR3 (PDB ID: 6S6C^30^), *Psp*R (PDB ID: 7W74^31^), *A*SR (PDB ID: 1XIO^32^), *Ns*XeR (PDB ID: 6EYU^33^), *Np*SRII (PDB ID: 1H68^34^), *Np*HR (PDB ID: 3A7K^35^), *Mr*HR (PDB ID: 6XL3^36^), *Hs*HR (PDB ID: 1E12^37^), BPR *Med*12 (PDB ID: 4JQ6^38^), ESR (PDB ID: 4HYJ^39^), XR (PDB ID: 3DDL^40^), GR (PDB ID: 6NWD^41^), TR (PDB ID: 5AZD^42^), NM-R3 (PDB ID: 5B2N^43^), KR2 (PDB ID: 3X3C^44^), VirR_DTS (PDB ID: 6JO0^45^), SzR4 (PDB ID: 7E4G^46^), ARI (PDB ID: 5AX0^47^), LR (PDB ID: 7BMH^48^), C1C2 (PDB ID: 3UG9^49^), ChR2 (PDB ID: 6EID^50^), Chrimson (PDB ID: 5ZIH^14^), ChRmine (PDB ID: 7W9W^51^), *Gt*ACR1 (PDB ID: 6CSM^52^), *Sr*Rh-PDE (PDB ID: 7CJ3^53^), HeR 48C12 (PDB ID: 6UH3^54^), *Ta*HeR (PDB ID: 6IS6^55^), *Cs*R (PDB ID: 6GYH^56^), *Tara*-RRB (PDB ID: 7PL9^17^). Inappropriate gaps in transmembrane helices were manually removed according to the 3-D structures.

The evolutionary history was inferred using the Neighbor-Joining method^57^. The optimal tree with the sum of branch length = 36.36238546 is shown. The nodes with the bootstrap values ≥ 80 % in the bootstrap test (1000 replicates) were indicated by black circles in Supplementary Fig. 458. The tree is drawn to scale, with branch lengths in the same units as those of the evolutionary distances used to infer the phylogenetic tree. The evolutionary distances were computed using the Poisson correction method^59^ and are in the units of the number of amino acid substitutions per site. The analysis involved 88 amino acid sequences. All positions containing gaps and missing data were eliminated. There were a total of 146 positions in the final dataset. Evolutionary analyses were conducted in MEGA6^60^.

### DNA constructs

Codon-optimized coding sequence of ChR024 was synthesized for expression in human cells and cloned into pCMV3.0-eYFP/mCherry vector between EcoRI and BamHI restriction sites. A Kir2.1 membrane trafficking signal, eYFP or mCherry, and an ER-export signal^61^ were fused to the C-terminus of ChR024. Other sequences of single mutants were similarly cloned into the pCMV3.0-eYFP/mCherry vector using the same strategy.

### Protein expression for hydroxylamine bleach experiment

Rhodopsins were expressed in COS-1 cells (JCRB9082), which were obtained from the Japanese Collection of Research Bioresources Cell Bank (Japan). COS-1 cells were transfected with a plasmid DNA using a polyethyleneimine-based method^62^, and all-*trans*-retinal was added at a final concentration of 2.5 μM, 4 hr post-transfection. Transfected cells were incubated at 37 °C with 5% CO_2_ for 48 hr. Rhodopsin-containing membranes of collected cells were solubilized with a phosphate buffer (100 mM NaCl, 50 mM phosphate, pH 8.0) containing 3% *n*-dodecyl-β-D-maltoside (DDM; Anatrace, OH) by shaking at 4 °C for 1 hr. After centrifugation (21,600 ×*g*, 4 °C, 10 min), the supernatant was collected for hydroxylamine bleach experiment.

For hydroxylamine bleach, solubilized rhodopsin samples were bleached by illumination with light (*λ* > 500 nm) from the output of a 1 kW Xe lamp (MAX-303, Asahi Spectra, Japan) through a long-pass filter (Y52, AGC Techno Glass, Japan) and a heat absorption filter (HAF50S15H, SIGMAKOKI, Japan) in the presence of 10 or 500 mM hydroxylamine (NH_2_OH). Absorption spectra were recorded before and after illumination using a UV-visible spectrophotometer (V-730, JASCO, Japan). Difference spectra were obtained by subtracting each post-illumination spectrum from the corresponding pre-illumination spectrum.

### Cell culture and transfection for electrophysiology

ND7/23 cells were grown in Dulbecco’s modified Eagle’s medium (D-MEM, FUJIFILM Wako Pure Chemical Co., Japan) supplemented with 5% fetal bovine serum (FBS) under a 5% CO_2_ atmosphere at 37 °C. ND7/23 cells were attached onto a collagen-coated 12-mm coverslips (4912-010, IWAKI, Japan) placed in a 4-well cell culture plate (30004, SPL Life Sciences, Korea). The expression plasmids were transiently transfected in ND7/23 cells using Lipofectamine^TM^ 3000 transfection reagent (Thermo Fisher Scientific Inc., MA). Six hours after the transfection, the medium was replaced by D-MEM containing 10% horse serum, 50 ng/mL nerve growth factor-7S (Sigma-Aldrich, MO), 1 mM N^6^,2’-O-dibutyryladenosine-3’,5’-cyclic monophosphate sodium salt (Nacalai tesque, Japan), and 1 μM Cytosine-1-β-D(+)-arabinofuranoside (FUJIFILM Wako Pure Chemical Co., Japan). Electrophysiological recordings were conducted at 3–7 days after the transfection. The transfected cells were identified by the presence of eYFP/mCherry fluorescence under an up-right microscope (BX50WI, Olympus, Japan).

### Electrophysiology

All experiments were carried out at room temperature (20–22 °C). Currents were recorded using an EPC-8 amplifier (HEKA Electronic, Germany) under a whole-cell patch clamp configuration. The data were filtered at 1 kHz, sampled at 50 kHz (Digidata1440 A/D, Molecular Devices Co., CA) and stored in a computer (pClamp11.1, Molecular Devices, CA). The standard internal pipette solutions for the whole-cell voltage clamp recordings from the ND7/23 cells contained 132 mM NaOH, 90 mM glutamate, 5 mM EGTA, 49 mM HEPES, 2.5 mM MgCl_2_, 2.5 mM MgATP, 2.5 μM ATR (pH 7.4 adjusted with HCl). The standard extracellular Tyrode’s solution contained 138 mM NaCl, 3 mM KCl, 2.5 mM CaCl_2_, 1 mM MgCl_2_, 10 mM HEPES, 4 mM NaOH, and 11 mM glucose (pH 7.4 adjusted with HCl). In the ion selectivity measurement, extracellular solution contained 146 mM XCl (X = Na^+^, K^+^, Li^+^, Cs^+^), 6 mM N-methyl-D-glucamine (NMG), 10 mM HEPES, 11 mM glucose (pH 7.4 adjusted with HCl). For NMG substitution, an extracellular solution was used containing: 146 mM NMG–HCl, 10 mM HEPES, and 11 mM glucose (pH 7.4 adjusted with NMG). For the divalent cation selectivity, bath solution contained 10 mM XCl_2_ (X = Ca^2+^, Mg^2+^), 131 mM NMG, 10 mM HEPES, 11 mM glucose (pH 7.4 adjusted with HCl). The pipette solution contained 130 mM NMG, 90 mM glutamate, 5 mM EGTA, 50 mM HEPES, 2.5 mM MgSO_4_, 2.5 mM MgATP, 2.5 μM ATR (pH 7.4 adjusted with H_2_SO_4_). For ion-selectivity experiments, after forming a gigaohm seal and establishing the whole-cell configuration in the extracellular Tyrode’s solution, the bath was exchanged for external solutions containing the indicated monovalent or divalent cations. Plasmid constructs of pCMV3.0-eYFP vector were used for these experiments.

For whole-cell voltage clamp, illumination at 390, 438, 475, 513, 549, 575 or 632 nm was carried out using a SpectraX light engine (Lumencor Inc., OR) controlled by pClamp 11.1 software. For the photocurrent detection, the power of the light at 575 nm was directly measured under a microscope using a visible light-sensing thermopile (MIR178 101Q, SSC Co., Ltd., Japan) and was 76.6 mW mm^−2^. The action spectrum was studied at a holding potential of −40 mV at wavelengths (nm, >90% of the maximum) with equivalent power density of 10.7 mW mm^−2^. Each action was estimated by the maximal amplitude of the photocurrent scaled by the light power density under the assumption of a linear relationship. In the action-spectrum measurement, ChR024 WT and P206T constructs were expressed by using pCMV3.0-mCherry vector. For Laser-flash patch clamp experiment, a laser flash (3–5 ns) at 532 nm (Nd:YAG laser, Minilite II, Continuum, CA) was used.

### Protein purification

The wild-type ChR024 (residues M1–T280) was modified to include a C-terminal Kir2.1 membrane-targeting sequence, a human rhinovirus 3C protease cleavage site, enhanced green fluorescent protein (eGFP), and an 8×His tag, and cloned into the pFastBac vector. Constructs were expressed in *Spodoptera frugiperda* (Sf9) insect cells using the pFastBac baculovirus system. Sf9 cells were grown in suspension to a density of 3.0 × 10^6^ cells ml^−1^, infected with baculovirus, and shaken at 27.5 °C for 24 hr. All-*trans*-retinal (ATR; FUJIFILM Wako Pure Chemical Co., Japan) was supplemented to a final concentration of 10 µM in the culture medium 24 hr after the infection. Cells were harvested and resuspended in a hypotonic lysis buffer (20 mM HEPES-NaOH pH 7.5, 20 mM NaCl, 10 mM MgCl_2_, 1 mM benzamidine, 1 µg ml^−1^ leupeptin, 10 µM ATR). The cell suspension was collected by centrifugation at 9,000 ×*g* for 10 min, and this process was repeated twice. The cell pellets were then disrupted using with a glass dounce homogenizer in a hypertonic lysis buffer (20 mM HEPES-NaOH pH 7.5, 1 M NaCl, 10 mM MgCl_2_, 1 mM benzamidine, 1 µg ml^−1^ leupeptin, 10 µM ATR), and the crude membranes were collected by ultracentrifugation (125,000 ×*g* for 1 hr) (Optima XE-90 and Type 45Ti rotor, Beckman Coulter, USA). The procedure was repeated twice, and the membranes were subsequently homogenized in membrane storage buffer (20 mM HEPES-NaOH pH 7.5, 500 mM NaCl, 10 mM imidazole, 20% (v/v) glycerol, 1 mM benzamidine, 1 µg ml^−1^ leupeptin), flash-frozen in liquid nitrogen, and stored at −80 °C until use.

For solubilization, the membrane fraction was incubated in a solubilization buffer (1% (w/v) *n*-dodecyl-β-D-maltoside (DDM) (Cat# D97002, Glycon Biochemicals, Germany), 20 mM HEPES-NaOH pH 7.5, 500 mM NaCl, 20% (v/v) glycerol, 10 mM imidazole, 1 mM benzamidine, 1 µg ml^−1^ leupeptin) at 4 °C for 2 hr. The insoluble cell debris was removed by ultracentrifugation (45Ti rotor, 125,000 ×*g* for 1 hr), and the supernatant was incubated with the Ni-NTA superflow resin (Cat# 30430, QIAGEN, Netherlands) at 4 °C for 1 hr. The Ni-NTA resin was loaded onto an open chromatography column, washed with 2.5 column volumes of wash buffer (0.05% (w/v) DDM, 20 mM HEPES-NaOH pH 7.5, 100 mM NaCl, and 50 mM imidazole) three times, and eluted with elution buffer (0.05% (w/v) DDM, 20 mM HEPES-NaOH pH 7.5, 100 mM NaCl, and 300 mM imidazole). After tag cleavage with His-tagged 3C protease, the sample was reapplied onto the Ni-NTA open column to remove the cleaved eGFP-His-tag and His-tagged 3C protease. The flow-through fraction was collected and concentrated to approximately 2 mg ml^−1^ using an Amicon Ultra centrifugal filter unit (50 kDa cutoff) (Merck Millipore). The concentrated samples were ultracentrifuged (71,680 ×*g* for 30 min) using TLA 55 rotor (Beckman Coulter, USA) before size-exclusion chromatography on a Superdex 200 Increase 10/300 GL column (Cytiva), equilibrated in SEC buffer (0.03% (w/v) DDM, 20 mM HEPES-NaOH pH 7.5, 100 mM NaCl). The peak fractions of the protein were collected and concentrated to approximately 5 mg ml^−1^.

### pH titration

To investigate the pH dependence of the absorption spectra of ChR024, concentrations of proteins were adjusted to OD = ∼0.25 at *λ*_max_ and solubilized in 6-mix buffer (10 mM trisodium citrate 10 mM MES, 10 mM HEPES, 10 mM MOPS, 10 mM CHES, 10 mM CAPS, 100 mM NaCl, 0.03 % DDM (pH 7.0)). The pH was adjusted to the desired value by adding small aliquots of HCl and NaOH. The absorption spectra were recorded with UV–vis spectrometer (V-750, JASCO, Japan). The measurements were carried out at every 0.3–0.6 pH.

### Laser flash photolysis

The experimental setup for laser flash photolysis measurement was similar to that reported previously^63,64^. The purified protein samples were solubilized in a solvent containing 20 mM HEPES (pH 7.5), 100 mM NaCl, 0.03% DDM. The absorption of ChR024 solution was adjusted to 0.5 at excitation wavelengths of 532 nm. The nanosecond second harmonic of a Nd-YAG laser (*λ* = 532 nm, INDI40, Spectra-Physics) was used for the excitation of rhodopsins to determine the transient absorption spectra and the time courses of the transient absorption change at specific probe wavelengths. The transient absorption spectra were obtained by monitoring the intensity change of white light from a Xe-arc lamp (L9289-01, Hamamatsu Photonics), passing through the sample, using an ICCD linear array detector (C8808-01, Hamamatsu Photonics, Japan). To improve the signal-to-noise (S/N) ratio, 60 spectra were averaged, and the singular value decomposition (SVD) analysis was performed. The time evolution of the transient absorption change was obtained by observing the intensity change of the output of a Xe arc lamp (L9289-01, Hamamatsu Photonics, Japan) monochromated by a monochromator (S-10, SOMA OPTICS, Japan) and passed through the sample after photoexcitation using a photomultiplier tube (R10699, Hamamatsu Photonics, Japan) equipped with a notch filter (532 nm, bandwidth = 17 nm) (Semrock, NY) to remove the scattered pump pulse. To enhance the S/N ratio, 100–200 signals were averaged, and the resulting data were recorded in a digital storage oscilloscope (DPO7104, Tektronix, OR). The signals were globally fitted with a multiexponential function to determine the lifetimes and absorption spectra of each photointermediate.

### High-performance liquid chromatography (HPLC) analysis of retinal isomers

Retinal configurations were analyzed by HPLC as described elsewhere^51^. Briefly, the chromophore retinal of the purified ChR024 was converted to retinal oxime by adding methanol and HA. Retinal oxime was extracted with *n*-hexane and injected into an HPLC system (pump: PU-4580; UV-visible detector: UV-4570, JASCO, Japan) with a silica column (particle size 3 μm, 150 × 6.0 mm; Pack SIL, YMC, Japan). Illumination was performed using a 1 kW Xe lamp (MAX-303) with a band-pass filter, HMX0590 (590 ± 5 nm; Asahi Spectra, Japan) for 1 min at room temperature. For light-adapted conditions, samples after illumination were kept for 10 s without (Light-adapted 1) or with (Light-adapted 2) 10 pulses of a nanosecond pulsed laser (532 nm, 4.2 mJ/pulse, 1.1 Hz). The molar compositions of retinal-oxime isomers were calculated from the peak areas and molar extinction coefficients at 360 nm (all-*trans*-15-*syn*: 54,900 M^−1^ cm^−1^; all-*trans*-15-*anti*: 51,600 M^−1^ cm^−1^; 13-*cis*-15-*syn*, 49,000 M^−1^ cm^−1^; 13-*cis*-15-*anti*: 52,100 M^−1^ cm^−1^; 11-*cis*-15-*syn*: 35,000 M^−1^ cm^−1^; 11-*cis*-15-*anti*: 29,600 M^−1^ cm^−1^; 9-*cis*-15-*syn*: 39,300 M^−1^ cm^−1^; 9-*cis*-15-*anti*: 30,600 M^−1^ cm^−1^).

## Reporting Summary

Further information on experimental design is available in the Nature Research Reporting Summary linked to this article.

## Data Availability

Data supporting the findings of this manuscript are available from the corresponding author upon reasonable request.

## Acknowledgments

This work was supported by Japan Science and Technology Agency (JST) CREST (grant No. JPMJCR1502 to I.T.; JPMJCR21P3/JPMJCR23B1 to H.E.K., JPMJCR22N2 to K.I.), JST PRESTO (JPMJPR24OF to M.F.), JST FOREST (JPMJFR204S to H.E.K.), Grants-in-Aid from the Japan Society for the Promotion of Science (JSPS) for Scientific Research (KAKENHI grant Nos. JP23H04863/JP25K09698 to T.N.; JP23K05007 to M.Ko.; JP24H02262 to M.F., JP19H03163/JP22H00400/JP25H01338 to H.E.K.; JP24H02268/JP25H00424 to K.I.), AMED (24bm1123057h0001 to H.E.K.), MEXT Promotion of Development of a Joint Usage/ Research System Project: Coalition of Universities for Research Excellence Program (CURE) (grant JPMXP1323015482 to K.I.), the Israel Science Foundation (Research Center grant 3131/20 to O.B.) and the European Commission, under Horizon Europe’s research and innovation programme (Bluetools project, Grant Agreement No. 101081957 to O.B.), and the Research Foundation for Opto-Science and Technology (to M.Ka.). K.I. and I.T. received support from RIKEN AIP; O.B. received support from the Louis and Lyra Richmond Memorial Chair in Life Sciences. The work conducted by the U.S. Department of Energy Joint Genome Institute (https://ror.org/04xm1d337), a DOE Office of Science User Facility, is supported by the Office of Science of the U.S. Department of Energy operated under Contract No. DE-AC02-05CH11231.

## Author contributions

S.T., C.Z., Y.K., T.N., M.Ko., N.M., H.Y., K.I. performed electrophysiological and spectroscopic characterization. M.Ka, Y.I. and I.T. constructed the machine learning model and predicted the *λ*_max_ of putative channelrhodopsin genes. F.S., O.B., and T.W. collected and inspected sequences of putative channelrhodopsin genes. M.W., M.F., and H.E.K. prepared purified protein samples. S.T., M.K., F.S., T.N., T.W., I.T., and K.I wrote the manuscript. All authors discussed and commented on the manuscript. H.E.K. supervised the research.

