## Supplementary Information for "Functional characterization of red-shifted rhodopsin channels from giant viruses explored by a machine-learning model for long-wavelength optogenetics"

Takaramoto et al.

**Supplementary Table 1. Twenty-three ChR-like genes that were predicted to have the most red-shifted $\lambda_{max}$ and expressed in COS-1 cells.**

| Amino-acid residues constituting the retinal binding pocket | Predicted $\lambda_{max}$ (nm) |
|---|---|
| SVIYSWTCIMIGFSCGWYPWHDAK (ChR099) | 548.2 |
| LISYEFGTLMMGFSCFWYPWHDAK (ChR024) | 543.4 |
| CTNYGWTCLILGFCAGWFPFYDAK (ChR014) | 543.1 |
| FMFFGWTCLMIGYSWCWFGFVDTK (ChR152) | 534.2 |
| IGFPEWTCLVLGFGSCWFPFWNAK | 533.6 |
| CMLYGWTCIMIGFASYWFPWHDSK (ChR255) | 527.7 |
| AAVYGWTCLMLAYGCCWYAWYDSK | 525.1 |
| FNMYQYTCIMLAFGAFWFPWFDAK | 524.5 |
| ELSYSWTCLMIGFAMGWFPWYDSK (ChR117) | 517.3 |
| SATYAWTCTTIGFGVGWFPFHDAK | 516.1 |
| FLCYEWTCMMLGFGVLWFPWLNAK | 515.3 |
| AAVYGWTCIMLAYGICWYAWYDSK | 512.7 |
| SALYEWTFLALGIAAVWYPWFDAK | 512.1 |
| MCVTLWTVIMIGFAMFWYPFFDSK (ChR264) | 510.4 |
| IGFPEWTCIMLGFGFWWFPFWNSK (ChR098) | 509.4 |
| SCVYEWTCLLTGFGCGWFPFHDAK | 508 |
| SSTYGWTCLLMGFAILWFGWFDSK | 506 |
| SSTFGWTCLLMGFSLLWFGWFDSK | 505.6 |
| CLKYEWTCICVGFGMGWYPFTDTK | 502.2 |
| CLHYAWSCITIGFGLGWFPFHESK | 501.9 |
| SSLYGWTCILIGFALGWYPFHDSK | 501.5 |
| AAVYEWTCITIGYGCGWFPFHDSK | 500.8 |
| IGFPEWTCMVLGYGLAWFPFWNSK | 500.6 |

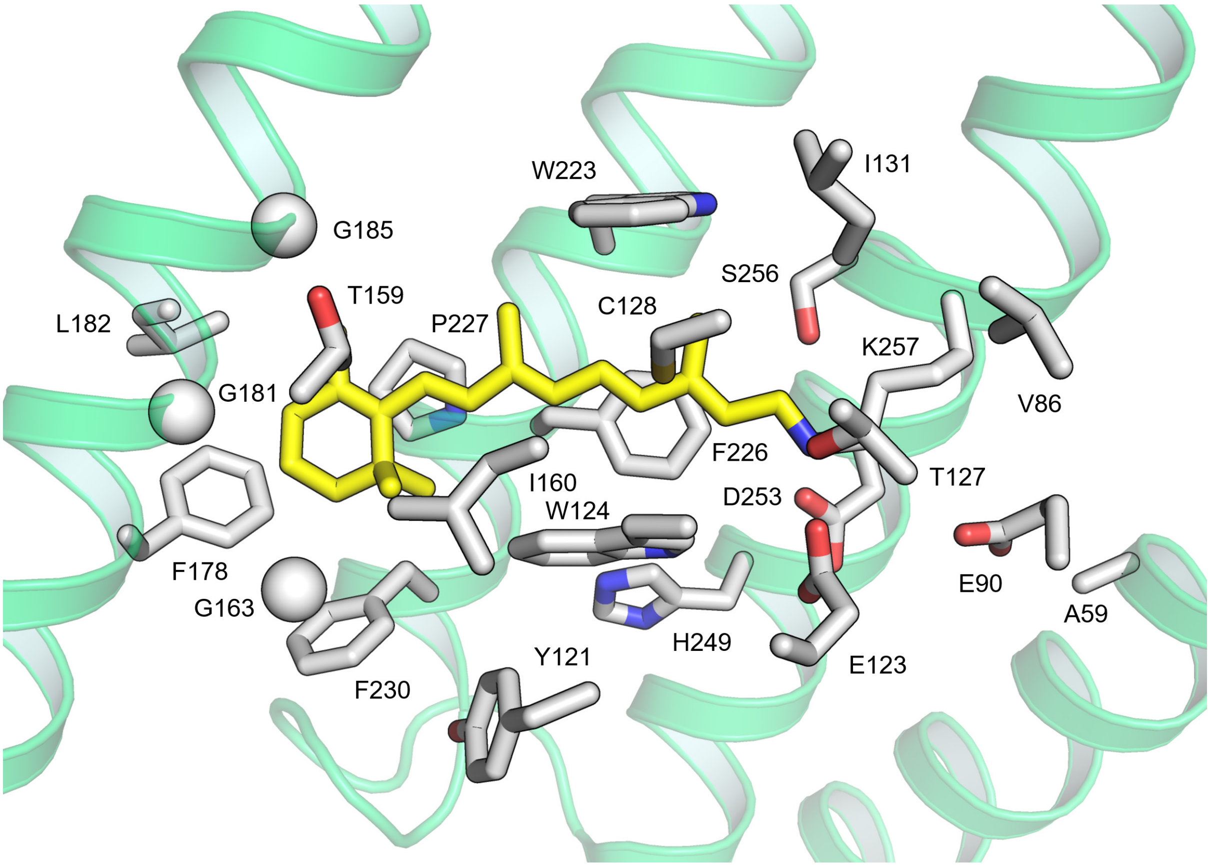


**Supplementary Fig. 1. Amino-acid residues constituting the retinal binding pocket used for the construction of the ML model and the $\lambda_{max}$ prediction.**

Twenty-four amino acid residues around the retinal chromophore that were known to play a critical role in determining $\lambda_{max}$ of microbial rhodopsins[9] in the X-ray crystallographic structure of channelrhodopsin 2 of *Chlamydomonas reinhardtii* (*Cr*ChR2, PDB ID: 6EID[50]). The corresponding residues were used for the construction of the ML model and the $\lambda_{max}$ prediction.

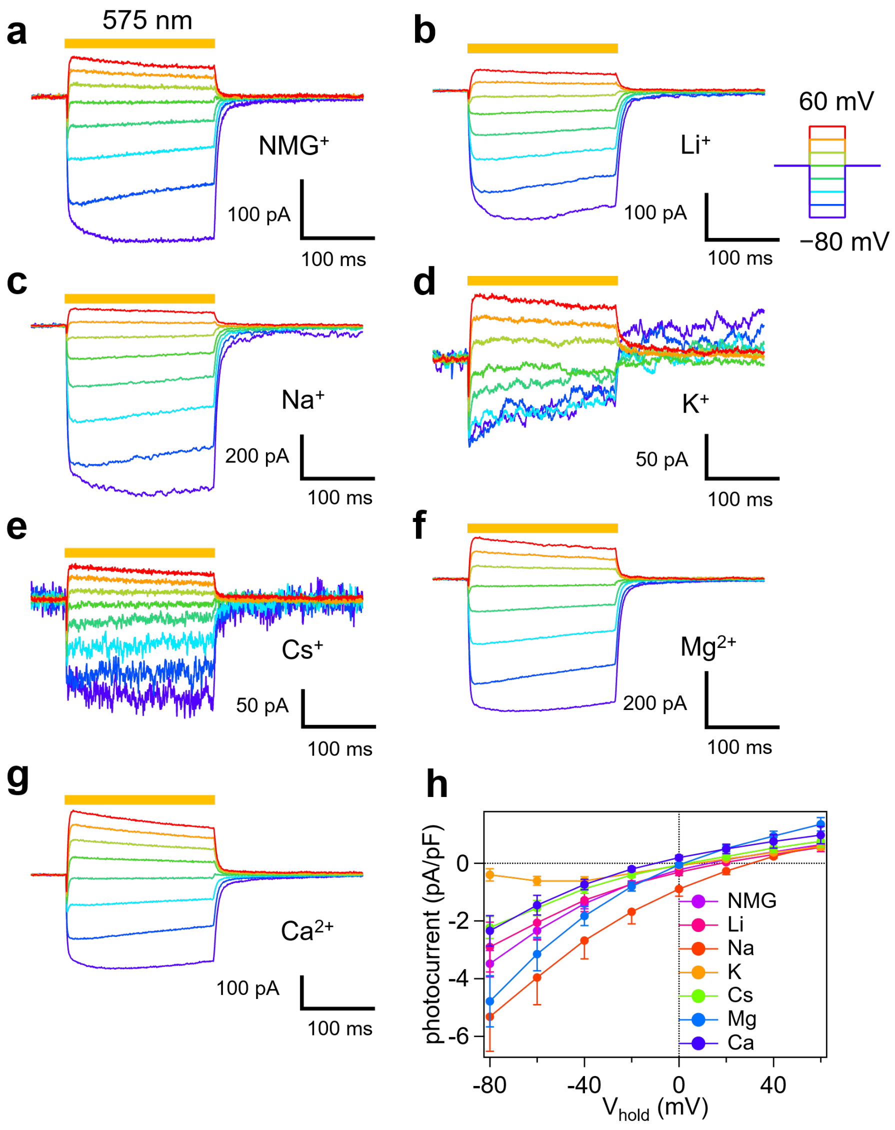


**Supplementary Fig. 2. Photocurrents of ChR024 in ND7/23 cells.**

**a–g** Photocurrents of ChR024 in ND7/23 cells with different cation species with 146 mM monovalent cations (NMG$^+$ (**a**), Li$^+$ (**b**), Na$^+$ (**c**), K$^+$ (**d**), and Cs$^+$ (**e**)) or 10 mM divalent cations (Mg$^{2+}$ (**f**) and Ca$^{2+}$ (**g**)) in the bath and 131 mM Na$^+$ in the pipette at pH$_{e/i}$ = 7.4. Holding potential was clamped from −80 mV to 60 mV by 20-mV steps in **a**. **h** *I–V* curves of the photocurrent of ChR024 with different cations in the bath solution.

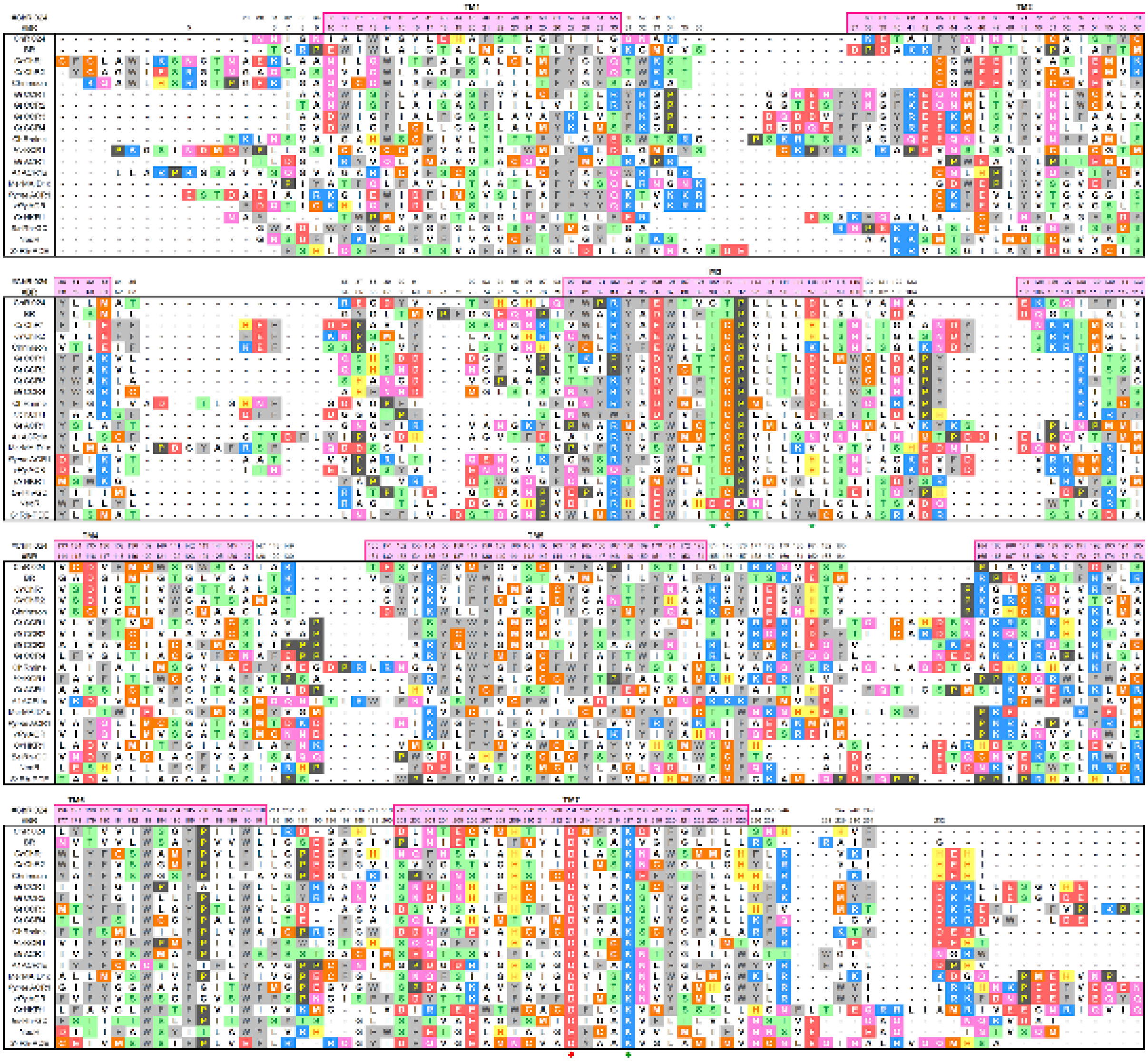

**Supplementary Fig. 3. Amino acid alignment of ChR024 with other microbial rhodopsins.**

The amino acid sequence of the transmembrane part of ChR024 is aligned with those of other microbial rhodopsins using the PROMALS3D server[25]. TMs in the X-ray crystallographic structure of BR (PDB ID: 1M0L[65]) and the motif residues, counterion in TM7, and retinal-binding lysine are indicated using green stars, a red rectangle, a green diamond. The amino acids are colored according to their chemical properties. Hydrophobic aliphatic residues: white, –OH group-bearing aliphatic residues: green, acidic residues: red, basic residues: blue, Asn and Gln: pink, sulfur-containing residues: orange; Pro: black; aromatic residues: gray.

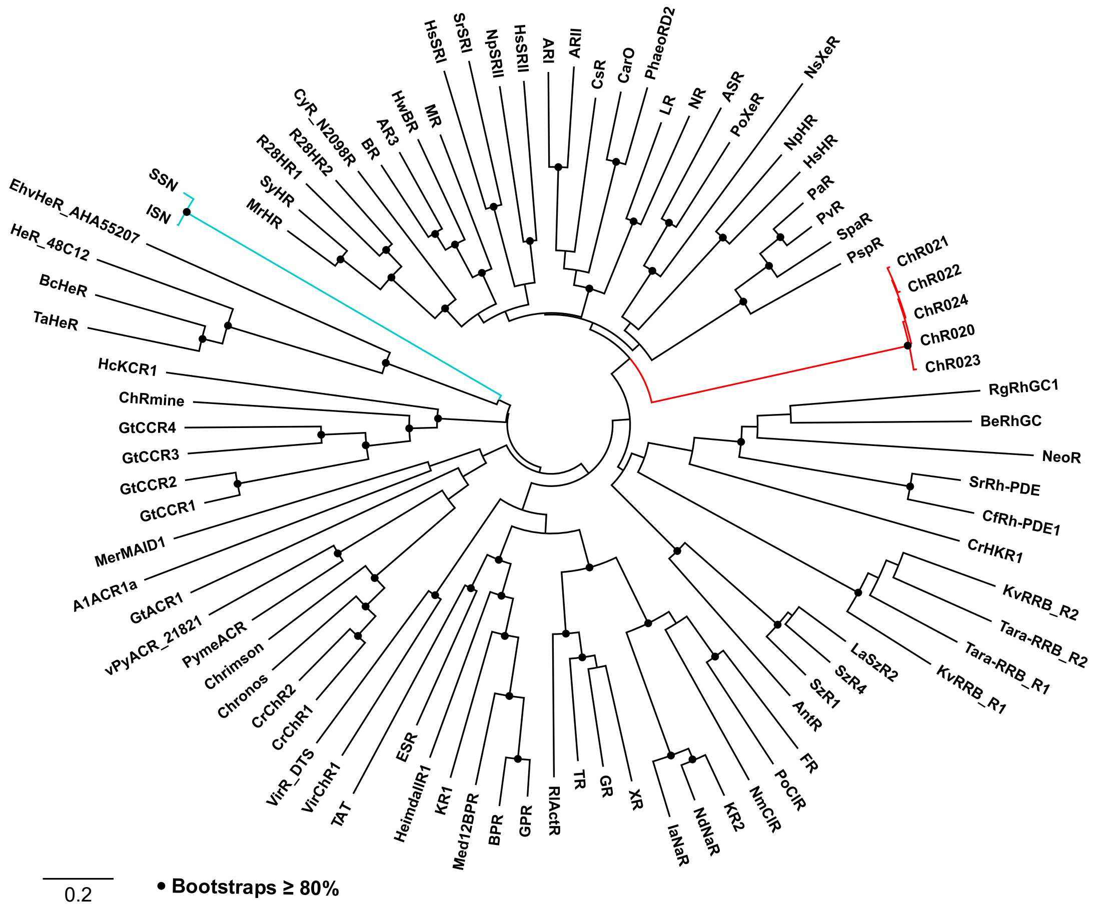


**Supplementary Fig. 4. The phylogenetic tree of microbial rhodopsins.**

The phylogenetic tree of microbial rhodopsin. The subgroups including ChR024 and SSN/ISN rhodopsins were highlighted in red and green lines, respectively. The nodes with bootstrap values ≥80% were indicated by black circles.

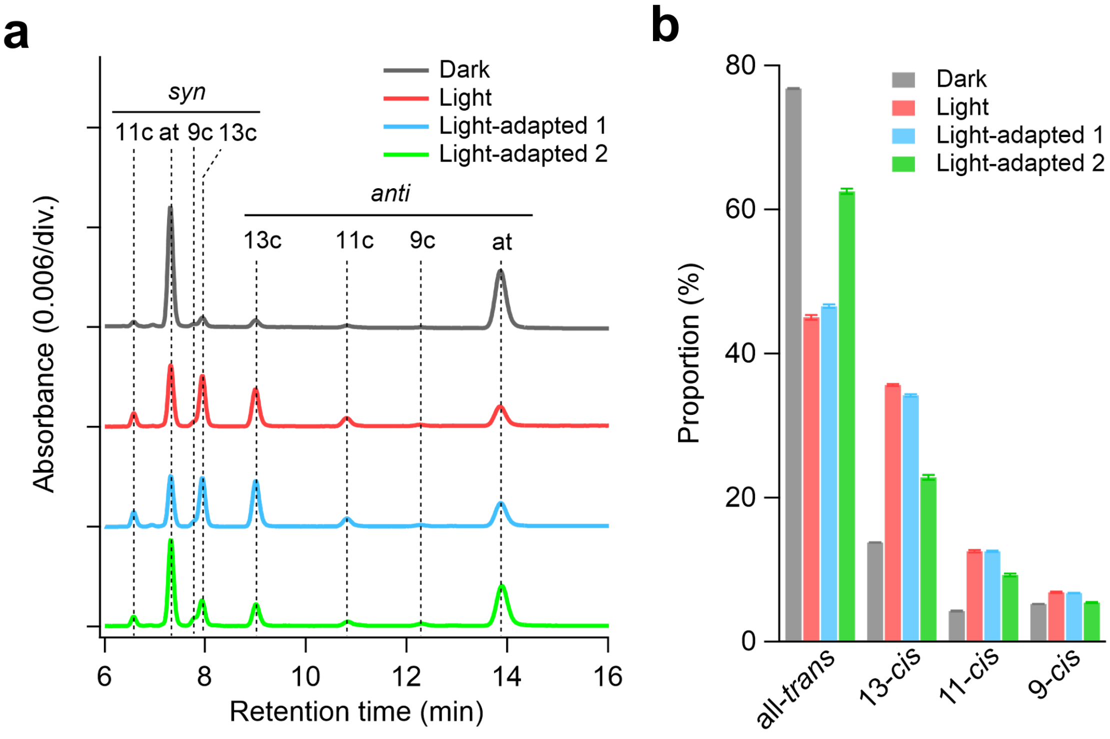


**Supplementary Fig. 5. HPLC analysis of retinal isomers in ChR024.**

**a** Chromatograms of retinal oximes derived from ChR024 under the dark-adapted condition (Dark), during illumination (Light), and under light-adapted conditions. For light adaptation, ChR024 samples underwent prolonged illumination, followed by either a 10-s dark incubation (Light-adapted 1) or exposure to 10 pulses of a 532-nm nanosecond pulsed laser (Light-adapted 2). 9c, 9-*cis*; 11c, 11-*cis*; 13c, 13-*cis*; at, all-*trans*. **b** Proportions of retinal isomers. Error bars indicate the standard error of the mean ($n$ = 3).

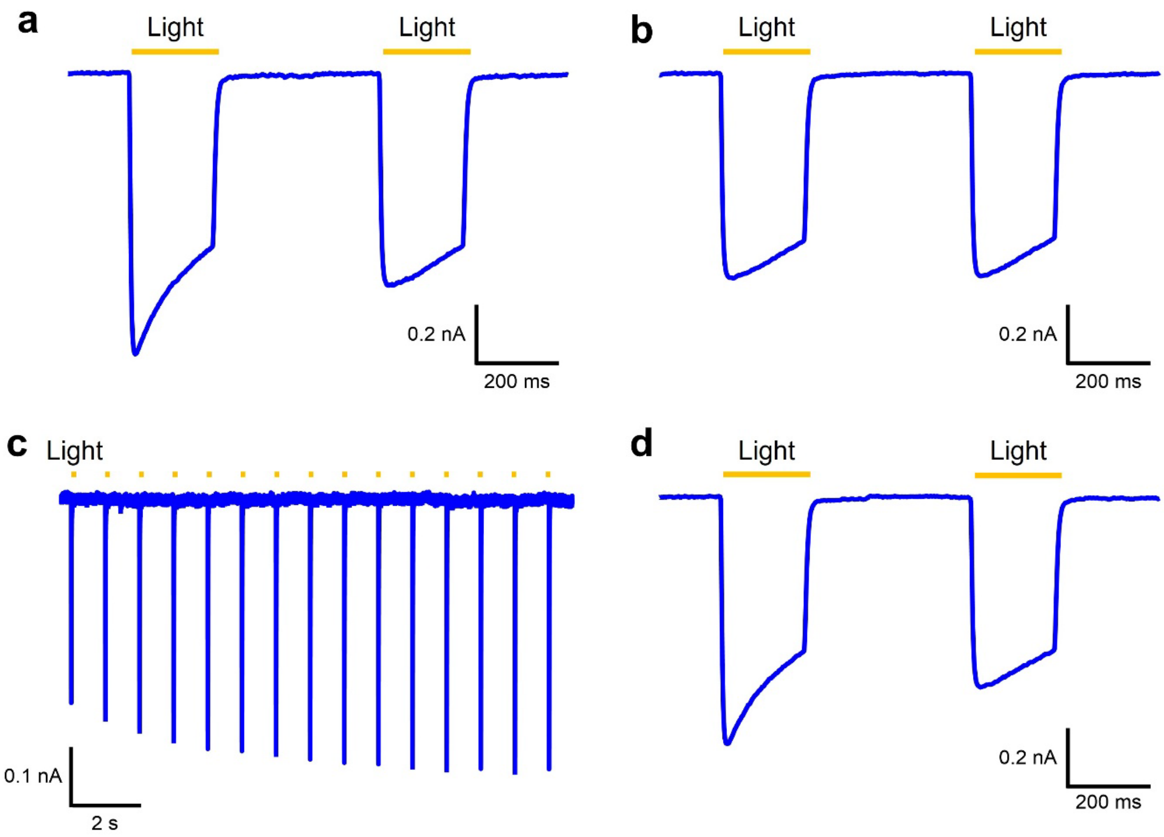


**Supplementary Fig. 6. Recovery of the peak current of ChR024.**

**a** The first recording of the photocurrent of ChR024 in the ND7/23 cells The cells were illuminated in the time regions (200 ms light pulse, 575-nm light) indicated by yellow lines. **b** Recording of photocurrents after ChR024 was deactivated during the recording of **a**. **c** Short light pulses to activate the ChR024 (2 ms light pulse, 575 nm yellow light) with 1-s interval between each light pulse. **d** Photocurrent of ChR024 with 200-ms light pulses after the 2 ms light pulses activation shown in **c.** All measurements were carried at −60 mV holding potential.

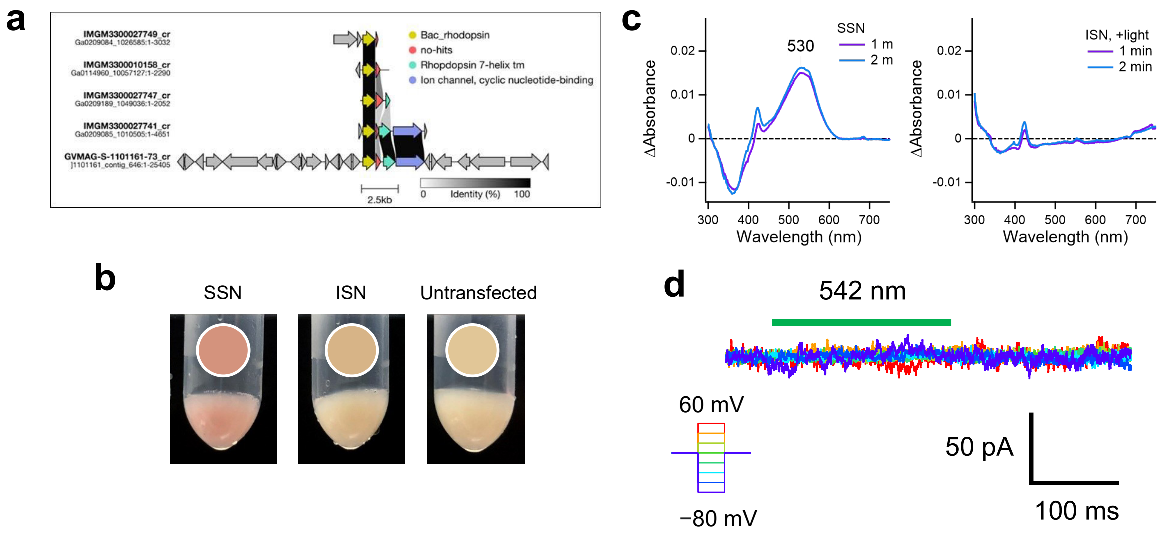


**Supplementary Fig. 7. SSN and ISN rhodopsin genes in the original scaffolds including ChR024**

**a** Genomic context of the scaffolds containing ChR024 and homologous genes. **b** Photographs of pellets of COS-1 cells transfected with SSN/ISN rhodopsin genes. **c** Difference absorption spectra of SSN rhodopsins in solubilized membrane, recorded before and after photobleaching via hydrolysis of RSB linkage in the presence of HA. **d** Photocurrents of SSN rhodopsin.
